# Effect of high proportion concentrate dietary on Yak jejunal structure, physiological function and protein composition during cold season

**DOI:** 10.1101/2020.10.19.345165

**Authors:** Jianlei Jia, Chunnian Liang, Xiaoyun Wu, Lin Xiong, Pengjia Bao, Qian Chen, Ping Yan

**Affiliations:** Key Laboratory of Yak Breeding Engineering Gansu Province, Lanzhou, Institute of Husbandry and Pharmaceutical Sciences, Chinese Academy of Agricultural Sciences, Lanzhou 730050, China; Key of laboratory of Plateau Ecology and Agriculture, Qinghai University, Xining 810016, China

**Keywords:** Feeding pattern (intensively and grazing), Morphology, Physiological function, Protein composition, Yak, Jejunum

## Abstract

The current study aimed to investigate the damage of long-term high concentrate diet feeding pattern on Yak jejunal structure, physiological function and protein composition during cold season. Twelve Datong male Yak (Bos grunniens) with the same age from cold season were randomly selected and slaughtered to determine Yak jejunal digestive enzyme activity, morphology and protein composition on different feeding patterns in Tibetan Plateau. The results showed that Yak jejunum digestive enzyme activity and morphology of grazing reared group were better than those in the intensively reared group. A total of 96 differentially expressed proteins were identified by label-free Mass Spectrometry (MS), which could be concluded to two predominant themes: protein structure and inflammatory response. Nine differentially expressed proteins were correlated in Yak jejunum damage in different feeding patterns. According to our study, feeding patterns resulted in the differences in Yak jejunum physiological function, morphology and protein composition. Our results confirmed that feeding of long-term high concentrate diet could damage the jejunum epithelial morphology and function.

## 1. Introduction

Yak (*Bos grunniens*) mainly lives above an altitude of 3000 meters in the Tibetan plateau and develops a dogged anti-adversity ability to exist, multiply and produce during the long period of natural selection artificial domestication ^[1]^. Yak production is an economic pillar of the Tibet plateau area. The Tibetan plateau is known for its extremely harsh conditions, characterized by high altitude, severe cold, low air oxygen, strong ultraviolet radiation and short forage growing season. Herbage and nutrients are insufficient to support the livestock during the cold period, especially when raised under traditional grazing management ^[2]^.

The modern intensive management pattern had gradually been replaced by traditional grazing patterns in the Tibet plateau area ^[3]^. The dietary concentration was more than 75% under modern intensive reared pattern, Yak bodyweight had a faster daily gain, and herdsman had a higher economic benefit. The ruminant needs an intake of 40%-70% dietary roughage to maintain normal rumen function and gastrointestinal microflora environment ^[4]^. However, long-term feeding of high concentrate diet can eventually have a serious impact on animal health, leading to rumen microorganisms’ metabolic disorders, such as accumulation of volatile fatty acids (VFAs) and decrease of pH decreased. Fermentable carbohydrate enters the small intestine through the rumen, causes intestinal acidosis, changes the microbial community structure and destroys the integrity of intestinal epithelial morphology and structure ^[5]^.

The dietary nutrients are digested in intestinal digestive enzymes and absorbed into the organism through intestinal epithelium to provide energy for animals ^[6]^. The jejunum is an important organ of nutrition digestion, absorption and metabolism, and a barrier against harmful substances. It plays an important role in Yak health. Jejunal congenital barriers can be damaged by external factors, leading to enteric flora disturbance and decreased production performance. Yak can graze on the pasture without dietary supplementation all the year in traditional grazing management, and the Yak body can firm a unique intestinal environment and digestive function to adapt to plateau grazing conditions. However, the modern intensive management pattern changes Yak diet habit, and high proportion concentrate diet reduces feedstuff residence time in Yak rumen and improves the amount of jejunum carbohydrate and jejunum microbial fermentation level, which has a negative effect on the monolayer structure of jejunum epithelial cells and damages the intestinal barrier ^[7,8]^. This may be an important physiological mechanism that a high concentrate diet destroys Yak jejunum structure and barrier. Here, we explore jejunal barrier dysfunction via long-term high concentrate dietary during cold season. Twelve Datong male Yak with different feeding patterns (intensively and grazing) during cold season were randomly selected and slaughtered, then determine Yak jejunal digestive enzyme activity with ELISA method and jejunal morphology with frozen section method. Using label-free method to identify proteins that specifically expressed with different feeding pattern, and we analyzed the signal pathway of target proteins. Collectively, our results shed new light on the damage of long-term high concentrate diet feeding on Yak jejunum. Meanwhile, this study laid a technical foundation in Yak intensively reared pattern.

## 2. Material and Methods

### 2.1 Ethics Statement

All procedures involving the use of animals were approved by the Animal Care Committee of Lanzhou Institute of Animal Science & Veterinary Pharmaceutics, Chinese Academy of Agricultural Sciences, China (QHDX-18-04-07-01). Animal slaughtering was approved by the National Administration of Cattle Slaughtering and Quarantine regulations (Qinghai, China).

### 2.2 Study Site

The study was performed at Haibei Tibetan Autonomous Prefecture, Qinghai Province, China, situated at northeastern Qinghai-Tibetan Plateau. This area is over 3000 m above sea level and has a dry cold climate.

### 2.3 Animals and Diets

Twelve Datong male Yaks (4±0.5 years old) were selected under grazing conditions on natural grasslands in Haibei Tibetan Autonomous Prefecture. All the Yaks were distributed into two groups. The Yaks of the experimental group were feed in a roofed shed via an intensively reared pattern, and the Yaks of the control group were feed in grazing pasture during the experimental period. The experiment lasted for 180 days from 16th November 2018 to 4th May 2019, and the first 10 d was an adjustment period of experimental group Yak. The following 170 d was for the data collection period.

Based on the determination of the feed intake of the grazing Datong Yak, the concentrated feed was prepared based on the recommendations for the Chinese Beef Cattle Raising Standard (NY/T 815-2004). The ingredients composition and nutritional level of Yaks of the experimental group are presented in Table 1. The roughage was a mixed ration of maize silage and oat hay in 1:1. The daily dietary of each Yak contained 4 kg concentrate and 2 kg roughage (DM) during the experimental period. The mixed diet nutrient composition was determined in the Qinghai University laboratory by ‘Feed Analysis and Quality Test Technology’ ^[9]^.

**Table 1.** Ingredients and proximate nutrition content of concentrate.

| Composition | Content (%) | Nutritional composition | Content |
| --- | --- | --- | --- |
| Maize | 67.0 | ME <sup>b</sup> | 3.01 Mcal/kg |
| Soybean meal | 5.0 | CP | 14.88 %DM |
| Cottonseed meal | 10.0 | Ca | 0.66 %DM |
| Wheat bran | 14.0 | P | 0.43 %DM |
| Salt | 0.5 |  |  |
| Sodium Bicarbonate | 1.5 |  |  |
| Stone Powder | 1.5 |  |  |
| Premix Feed <sup>a</sup> | 0.5 |  |  |

<sup>a</sup> The premix provided the following per kg of diets : VA 12000IU, VD 20 00IU, VE 30IU, Cu 12mg, Fe
64mg, Mn 56mg, Zn 60mg, I 1.2mg, Se 0.4mg, Co 0.4mg;
<sup>b</sup> ME was calculation value (NRC,2007), others were measured value.

### 2.4 Measurement of Samples

Datong male Yaks were stunned by the captive bolt pistol, and the blood was drained at the end of the experiment. The jejunum was immediately isolated after slaughtering, and jejunal contents (10 g) were aliquoted into 2 mL sterile tubes and then conserved at −80°C for digestive enzyme activity analysis. The activity of amylases (AMS), trypsinases (TRS), lipases (LPS) and carboxymethyl cellulose (CLS) were determined According to Fan’s method ^[10]^ using commercially available TaKaRa kits.

The separated jejunum tissue was divided into two parts. One part of jejunum tissue was fixed in 4% buffered formaldehyde for 72 hours. The samples were dehydrated under different concentrations of glucose and embedded in a frozen embedding medium. Tissue sections (5-7 µm) were cut and stained with hematoxylin and eosin (HE), displaying villous height, villous width, crypt depth, mucosa thickness and muscular thickness, respectively. Images were acquired using the DP70 software (Olympus, Nagano, Japan) per transverse section of jejunum tissue. The slides were observed using BX51 microscopy (Olympus, Nagano, Japan). Measurements were made using Image Proplus5.1 software. Ten well-oriented and intact crypt-villus units of each slide were measured in triplicate ^[11]^.

Another part was aliquoted into a 1.5 mL centrifugal tube when the jejunum surface contaminants were removed by washing with phosphate-buffered saline solution and conserved at −80 °C ultracold storage freezer for proteomics and RT-PCR analysis. Proteomics analysis was conducted by Novogene (China, Beijing). There were 3 replicates in each group. Jejunum proteins were extracted by lysis buffer method ^[12]^. Digestion of the protein (250 μg for each sample) was performed according to the FASP procedure. Label-free Mass Spectrometry (MS) experiments were performed on a Q Exactive mass spectrometer that was coupled to an Easy nLC system (Thermo Fisher Scientific, MA, USA). The MS data were analyzed using MaxQuant software and compared with the UniProt Bos taurus database. Relative quantitative real-time PCR (RT-PCR) was performed to determine the copy number of target genes. The primers were synthesized at Shanghai Biological Engineering Ltd., China. The RT-PCR reaction was carried out in a Light Cycler 480 fluorescence quantitative meter (Fritz Hoffmann-La Roche Co. Ltd., Basel, Switzerland).

### 2.5 Statistical Analysis

Bioinformatics was analyzed via R language toolkit by Novogene (China, Beijing). Functional annotation and classification of all identified proteins were determined using the Blast2GO and InterProScan program against the uniport database. Pathway analyses were extracted using the search pathway tool of the KEGG mapper platform (http://www.genome.jp/kegg/mapper.html) and BLAST program. Pathway enrichment statistics were conducted by the Fisher’s exact test, and the pathways with a corrected P > 0.05 were defined as the most significant pathways. The STRING program (http://string-db.org/) for the retrieval of interacting genes/proteins database for the prediction of the physical and functional interactions was used to analyze the PPIs. The graphical visualization and analysis of the interaction network were performed in Cytoscape software.

The data were expressed as mean ± standard deviation (SD). Duncan’s post hoc test was used to determine any significant differences between each group. Differences were considered significant at *P* < 0.05 and extremely significant at *P*<0.1.

## 3. Results

### 3.1 Digestive enzyme activity analysis of jejunum

The activity of AMS, LPS, TRS and CLS in different feeding patterns are presented in Table 2. The data indicated that the AMS and CLS activities of the grazing reared group was higher than that in the intensively reared group, but there was no significant difference between the two groups (*P*>0.05); The LPS activity of the intensively reared group was significantly greater than that in grazing reared group (*P*<0.05). The TRS activity of the intensively reared group was significantly lower than that in the grazing reared group (*P*<0.05). The Yak jejunum digestive enzyme activity of the grazing reared pattern was higher than the intensively reared pattern.

**Table 2.**
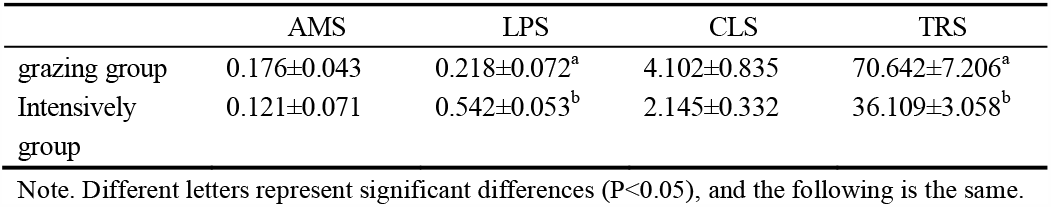
The alkaline phosphatase activity of yak jejunum in different feeding pattern.

### 3.2 Histological analysis of jejunum morphology

The villous height, villous width, crypt depth, mucosa thickness and muscular thickness in grazing reared and intensively reared groups are summarized in Table 3. The villous height, villous width, mucosa thickness and muscular thickness of jejunum were significantly greater in the grazing reared group than those in the intensively reared group (*P*<0.05). However, the control group’s crypt depth was lower than that in the intensively reared group, but no significant difference was observed between those two groups (*P*>0.05).

**Table 3.** The villous height, villous width, crypt depth, mucosa thickness and muscular thickness of yak jejunum in different feeding pattern.

|  | villous height | villous width | crypt depth | mucosa<br>thickness | muscular<br>thickness |
| --- | --- | --- | --- | --- | --- |
| grazing<br>group | 563.33±11.48<sup>a</sup> | 149.29±2.67<sup>a</sup> | 178.84±6.59 | 808.38±15.52<sup>a</sup> | 528.91±20.93<sup>a</sup> |
| intensively<br>group | 408.31±10.46<sup>b</sup> | 82.88±5.45<sup>b</sup> | 194.61±8.37 | 939.18±14.14<sup>b</sup> | 751.51±15.14<sup>b</sup> |

Note. Different letters represent significant differences (P<0.05), and the following is the same.

### 3.3 Changes in proteome profiles during different feeding pattern

We successfully identified 1208 proteins with Bos taurus database matching of Mascot and label-free method. Then we applied a manual thresholding approach and a probabilistic prediction algorithm, yielding 1066 high-confidence candidates. Of these 1066 proteins, 96 differentially expressed proteins were identified between the intensively reared group and grazing reared group using P < 0.05 or a quantitative ratio of > 1.2 or < 0.83 (Table S1). There were 17 up-regulated and 79 down-regulated differentially expressed proteins in the grazing reared group compared to that in the intensively reared group (Fig. 1).

**Fig 1.**
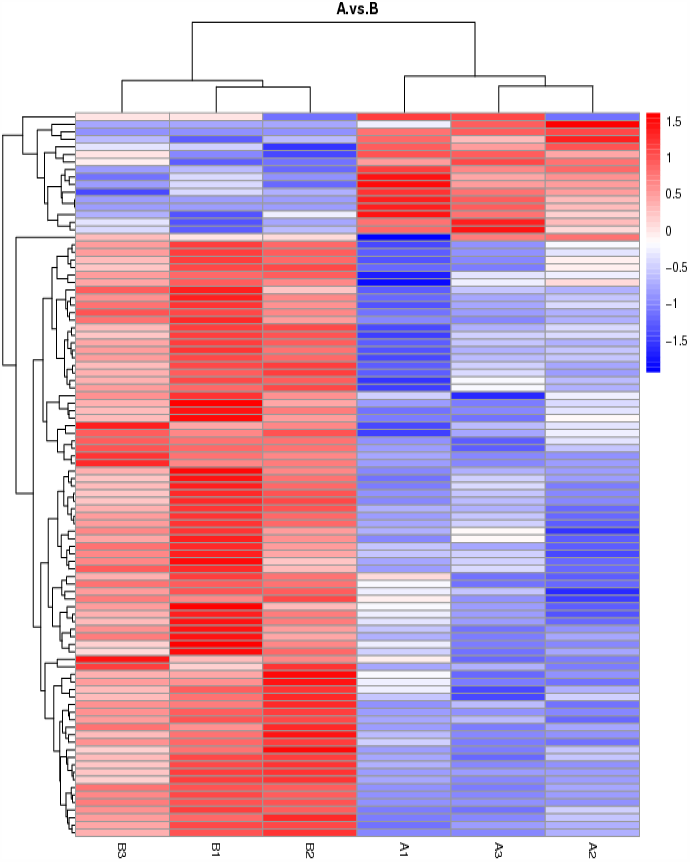
Differentially expressed proteins, including 3 replicates with yak jejunum during grazing reared group (A) and intensively reared group (B). The image presents the relative abundance of proteins using different colors, where deeper red represents higher intensity and blue represents lower intensity

### 3.4 Bioinformatics analyses

To explore biological functions associated with differentially expressed proteins of Yak jejunum during different feeding patterns, enrichment analysis in the Gene Ontology (GO) containing cellular components (CC), molecular function (MF) and biological process (BP) was performed (Fig. 2). The major enriched terms of differentially expressed proteins were membrane (GO:0016020), cytoplasm (GO:0005737) and mitochondrion (GO: 0005739; GO:0005740) for cellular components. Other enriched terms were molecular function in oxidoreductase activity (GO:0016491), cell-cell adherens junction (GO:0005525), adenylate kinase activity (GO:0004017), protein binding (GO:0019904) and structural molecule activity (GO:0003954). Apoptosis (GO: 0048468), inflammatory response (GO:0016020), cellular response to fatty acid (GO:0008152), actin cytoskeleton reorganization (GO:0055114) and adenylate cyclase-inhibiting G protein-coupled receptor (GO:0046034), which were enriched in biological processes.

**Fig 2.**
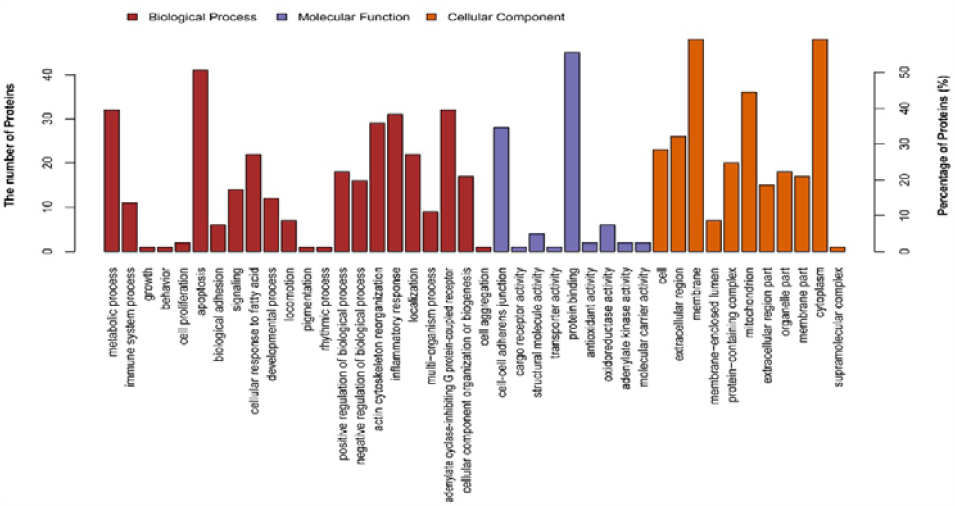
Enrichment GO terms Analysis.

The major pathways associated with differentially expressed proteins of Yak jejunum during grazing reared group and intensively reared group were identified using KEGG pathway analysis via KAAS software. The KEGG results showed that out of a total of 71 pathways, 49 were significantly enriched (p-value lower than 0.05).The top 6 of 20 significantly enriched pathways, ranked by significance and percent overlap are tight junction signaling pathway (ko04530), NF-kappa B signaling pathway (ko04064), MAPK signaling pathway (ko04010), cytokine-cytokine receptor interaction (ko04060), calcium signaling pathway (ko04020) and signaling pathways regulating pluripotency of stem cells (ko04550, Fig. 3). The results of the KEGG pathway analysis matched the results of the GO analysis.

**Fig 3.**
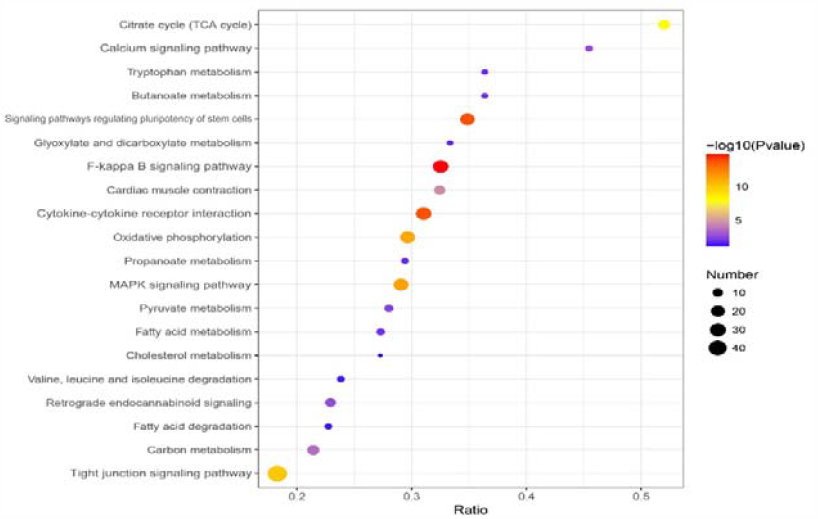
Enrichment KEGG pathways Analysis (Fisher’s Exact Test, P value < 0.01)

To gain information on interaction network, possible interactions among the 96 proteins were analyzed online using the Uniport database. As expected, the 96 target proteins constituted a complex and strong PPi network, and those results of this analysis identified two predominant themes: protein structure and inflammatory response. To provide further insights into the biological processes identified by this approach, we took Fishers’ test (Significance A/B Test) for target terms, those terms with the highest number of proteins from our results show the proteins that result in the identification of an enriched biological process. For example, the identification of the “Tight junction” (Occludin, Claudin, Zo-1, ATP1B3 and HKDC1) and “Inflammatory reaction” (GPR41, IL-6, TNF-α and CCL5) terms results in our list of differentially abundant proteins (Fig. 4).

**Fig 4.**
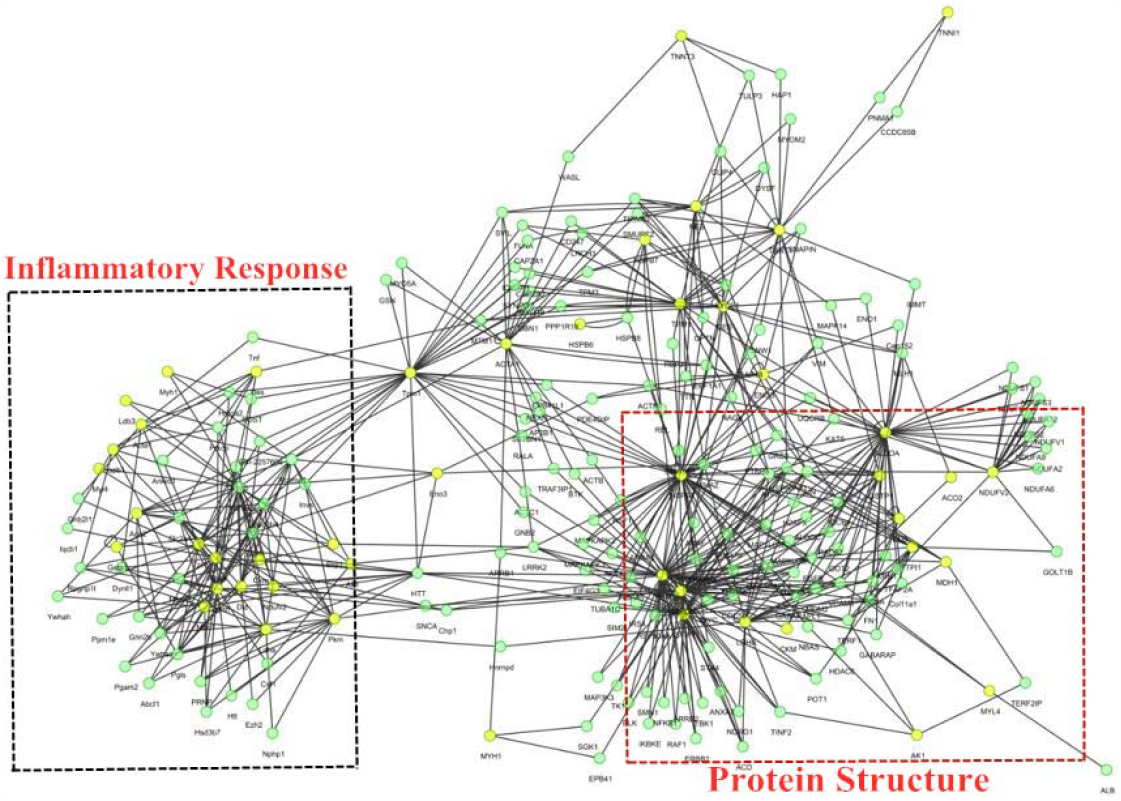
Protein-protein interaction networks of the differential abundance proteins of different feeding groups in Yak jejunum based on Cytoscape software. The nodes were proteins from Bos taurus database and the lines were the predicted functional annotations

### 3.5 Validation of differentially expressed proteins coding genes by RT-PCR

As shown in Fig. 5, the gene expressions of OCLN, CLDN1, TJP1 and HKDC1 were higher in the grazing reared group than those in the intensively reared group (*P*<0.05) via RT-PCR, and the ATP1B3 expression had no significant difference between the two groups. However, the gene expressions of IL6, GPR41, CCL5 and TNF were higher in the intensively reared group than those in the grazing reared group (*P*<0.05). In summary, the results of selected differentially expressed proteins coding genes by RT-PCR were the same expression tendency of the label-free analysis. The RT-PCR assay illustrated that the label-free results were reliable for further analyses.

**Fig 5.**
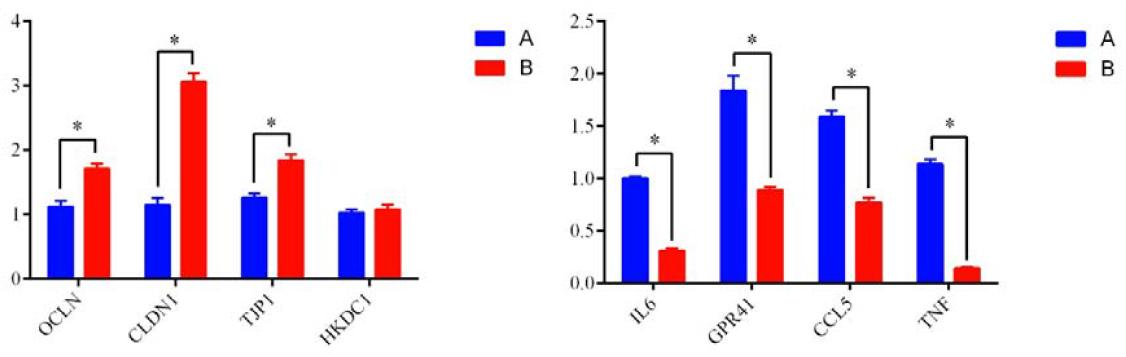
Effects of different feeding pattern on the differentially expressed proteins coding genes in yak jejunum during intensively reared group (A) and grazing reared group (B).

## 4. Discussion

Jejunum is the main organ for dietary nutrition digestion, absorption and metabolism, and plays an important role in Yak health. The dietary concentrate is more than 75% under modern intensive management pattern to enhance daily gain and performance. However, long-term high concentrate diet feeding has a serious impact on animal health, such as jejunal acidosis ^[4]^. The level of dietary nutrition digestion, absorption and metabolism can be reflected through digestive enzyme activity. Starch is the main energy source for ruminants. AMS can hydrolyze starch to glucose, maltose and oligosaccharides, which acts on the 4-c-D-glycosidic bonds of the starch molecule ^[13]^. Fat is the necessary material for energy storage and nutrition in the animal body, and the animal essential fatty acids are mainly provided by dietary fat. LPS can digest dietary fat to absorbable nutrients, such as free fatty acid, glycerin and monoglyceride. TRS can hydrolyze dietary proteins to small molecular amino acids, and those amino acids are involved in the pathway of intracellular signaling transmission and apoptosis. It plays a vital role in animal immune response ^[14]^. The ruminant cellulose-degrading is mainly dependent on the cellulase secreted by microorganisms, and CLS activity is an important ability to degrade cellulose ^[15]^. In our study, Yak jejunum digestive enzyme activity can be in a weakened state with long-term high concentrate diet feeding. We observed that the digestion and absorption of grazing reared patterns were better than the intensively reared group.

The morphophysiological variations in the jejunal epithelium of ruminants identified the ability of dietary nutrients digestion and absorption ^[11]^. Changes in the structure of villous absorptive surface (height, width and crypt) in jejunal epithelial were closely related to the extent of enterocyte development, which were direct representations of the quality of the intestinal environment as the indicator of intestinal health ^[16]^. The present study showed that the ability of dietary nutrients digestion and absorption increased with the increase of villous absorptive surface, and submucosa and muscular thickness of jejunum were decreased with long-term high concentrate diet, decreasing the absorption of nutrients in Yak jejunum. When the high concentrate diet was fed for a long-term, many nutrients were degraded into short-chain fatty acids (SCFAs) by microorganisms in the rumen, leading to an increase in the jejunum VFAs and a decrease in the jejunum pH value. VFAs had a strong lipid solubility when the pH value dropped below the critical value. VFAs can infiltrate into jejunum mucosa cells to acidify cells, damaging the intestinal barrier and causes an inflammatory response ^[17,18]^. We found that the significantly decreased resulted in the intensively reared group of Yak jejunum, such as villous height, villous width, mucosa thickness and muscular thickness. These results agreed with previous studies in ruminants.

SCFAs in rumen permeate into Yak jejunum, destroy the structural integrity of morphophysiological, and damage epithelial barrier function ^[19]^. Jejunal mechanical barrier plays a protective role in jejunum via the normal morphological structure of jejunal mucosa ^[20-22]^. Tight junction proteins are the important parts of the jejunum mechanical barrier composed of the transporter protein family, including occludin and claudin, which are mainly involved in enterocytes proliferation, differentiation, apoptosis and cell bypass permeability regulation. The damaged mechanical barrier induced jejunal dysfunction causes pathogens to infect enterocytes, which results in tight junction structural damage, permeability increasing and inflammatory response ^[19]^. According to recent reports and our study, target proteins can be concluded to two predominant themes: protein structure (tight junction signaling pathway, calcium signaling pathway and signaling pathways regulating pluripotency of stem cells) and inflammatory response (MAPK signaling pathway, chemokine signaling pathway and cytokine-cytokine receptor interaction). GPR 41, a key activator of the jejunal mechanical barrier, participates in regulating body inflammatory response and affects the tight junction signaling pathway ^[23]^. The large concentrate intake can start enterocytes gluconeogenesis process, activates the tight junction signaling pathway and regulates downstream related cell apoptosis pathways, decreasing the tight junction proteins expressions, such as occludin, claudin and Zo-1. Meanwhile, the proliferation of regulatory T cells promotes the inflammatory response pathway (MAPK) related gene expression to secrete many anti-inflammatory cytokines (IL-6, TNF-α and CCL5) ^[24-26]^. GPR41 was detected in the current study. Long-term high concentrate diet feeding changes the normal jejunal metabolic pathways. Several identified candidate proteins were identified by label-free MS, followed by qPCR, including occludin, claudin, Zo-1, ATP1B3, HKDC1, IL-6, TNF-α and CCL5. Compared with the grazing reared group, the expressing quantity of occludin, claudin, Zo-1, ATP1B3 and HKDC1 in Yak jejunum presented significantly down-regulated in the intensively reared group. However, IL-6, TNF-α and CCL5 showed a contrary tendency, indicating that many SCFAs in the jejunum of the high-concentrate diet could damage the mechanical barrier and reduce the differentially expressed proteins. At the same time, GPR41 was activated to up-regulate the expression and promoted the expression of the proinflammatory factors. In summary, the results provide evidence of the validity of the damage of the high concentrate diet on Yak jejunum.

## 5. Conclusions

We found that feeding pattern results in Yak different jejunum physiological function, morphology and protein composition. Long-term high concentrate diet feeding can damage the jejunum epithelial morphology and structure. Therefore, we suggest that the high concentrate diet is suitable for the short-term fattening pattern of reserved Yak.

## Supporting information

Certification for English editing

## Acknowledgments

The authors are grateful to all the participants who took part in this study. This research was funded by the Agricultural Science and Technology Innovation Program (CAAS-ASTIP-2014-LIHPS-01), the National Beef Cattle Industry Technology & System (CARS-38), National Natural Science Foundation of China (31972561).

## Abbreviations

ME: metabolic energy
CP: crude protein
Ca: Calcium
P: Phosphorus
DM: dry matter
AMS: amylases
LPS: lipases
CLS: carboxymethyl cellulase
CLS: carboxymethyl cellulase

