## Supplementary material for "Effect of high proportion concentrate dietary on Yak jejunal structure, physiological function and protein composition during cold season": Certification for English editing

MogoEdit

### CERTIFICATE OF ENGLISH EDITING

This is to certify that the manuscript entitled

commissioned to us has been carefully edited by a native English-speaking editor of MogoEdit, and the grammar, spelling, and punctuation have been verified and corrected where needed. Based on this review, we believe that the language in this paper meets academic journal requirements. Please contact us with any questions.

MOGOEDIT
MOGO INTERNET TECHNOLOGY CO., LTD.

*Gang Zhang*

Dr. Gang Zhang
Founder & CEO of MogoEdit

Date of Issue
December 22, 2020

**Disclaimer:** The changes in the document may be accepted or rejected by the authors in their sole discretion after our editing. However, MogoEdit is not responsible for revisions made to the document after our edit on **December 22, 2020**.

MogoEdit is a professional English editing company who provides English language editing, translation, and publication support services to individuals and corporate customers worldwide. As a company invested by the affiliate fund of Chinese Academy of Science, MogoEdit is one of the leading language editing service providers in China, whose clients come from more than 1000 universities and research institutes.
MogoEdit Website: http://en.mogoedit.com/
500+ native English editors: http://en.mogoedit.com/editors

Mogo make the world flatter
